# Ontogenetic Oxycodone Exposure Affects Early-Life Communicative Behaviors, Sensorimotor Reflexes, and Weight Trajectory in Mice

**DOI:** 10.1101/2020.09.30.321372

**Authors:** Elena Minakova, Simona Sarafinovska, Marwa O. Mikati, Kia Barclay, Katherine B. McCullough, Joseph D. Dougherty, Ream Al-Hasani, Susan E. Maloney

**Author notes:** Authors contributed equally (Elena Minakova, Simona Sarafinovska, Ream Al-Hasani, Susan E. Maloney). Correspondence: Dr. Ream Al-Hasani, Center for Clinical Pharmacology, St. Louis College of Pharmacy, Washington University School of Medicine, Campus Box 8054, 660 South Euclid Avenue, St. Louis, MO 63110-1093, Dr. Susan E. Maloney, Washington University School of Medicine, Department of Psychiatry, Campus Box 8232, 660 South Euclid Avenue, St. Louis, MO 63110-1093.

## Abstract

Nation-wide, opioid misuse among pregnant women has risen 4-fold from 1999 to 2014, with commensurate increase in neonates hospitalized for Neonatal Abstinence Syndrome (NAS). NAS occurs when a fetus exposed to opioids *in utero* goes into rapid withdrawal after birth. NAS treatment via continued postnatal opioid exposure has been suggested to worsen neurodevelopmental outcomes. We developed a novel model to characterize the impact of *in utero* and postnatal oxycodone (Oxy) exposure on early behavior and development. Via subcutaneous pump implanted before breeding, C57BL/6J dams were infused with oxycodone at 10 mg/kg/day from conception through pup-weaning. At birth, *in utero* oxy-exposed pups were either cross-fostered (paired with non-oxy exposed dams) to model opioid abstinence (short-oxy) or reared by their biological dams still receiving Oxy to model continued postnatal opioid exposure (long-oxy). Offspring from vehicle-exposed dams served as cross-fostered (short-veh) or biologically-reared (long-veh) controls. Short-oxy exposure resulted in sex-dependent weight reductions and altered spectrotemporal features of isolation-induced ultrasonic vocalization (USV). Meanwhile, long-oxy pups exhibited reduced weight and sex-differential delays in righting reflex. Specifically, long-oxy female offspring exhibited increased latency to righting reflex. Long-oxy pups also showed decreases in number of USV calls, and changes to spectrotemporal USV features. Overall, ontogenetic Oxy exposure was associated with impaired attainment of gross and sensorimotor milestones, as well as alterations in communication and affective behaviors, indicating a need for therapeutic interventions. The model developed here will enable studies of withdrawal physiology and opioid-mediated mechanisms underlying these neurodevelopmental deficits.

## INTRODUCTION

In the past two decades, illicit drug use and prescription opioid use in the United States has risen to epidemic proportions, with the US Department of Health declaring a public health emergency in 2017. The US Department of Health states that 46,802 people died from opioid overdose in 2018 and an estimated 2 million people have an opioid use disorder^1^. The public health crisis is largely driven by increased misuse of the prescription opioids hydrocodone, oxycodone (Oxy), and methadone^2,3^.

As a result of the opioid epidemic, the national prevalence of opioid use disorder among pregnant women has increased more than 4-fold from 1999 through 2014^2^. Consequently, there has been a significant increase in the number of neonates hospitalized for Neonatal Abstinence Syndrome (NAS), a constellation of withdrawal symptoms affecting the nervous system, gastrointestinal tract, and respiratory system following *in utero* exposure to opioids^4^. Currently, medical management of NAS involves keeping the infant swaddled in a low-stimulation environment with promotion of maternal-infant bonding^4^. In cases of moderate-severe NAS, neonatal withdrawal is managed by opioid replacement therapy to alleviate withdrawal symptomatology^4^. Overall, clinical studies have not addressed whether long-term neurobehavioral outcomes are improved by managing withdrawal, or whether continued postnatal exposure to opioids and adjunct agents used for withdrawal management worsen long-term outcomes^5^.

Epidemiological evidence suggests *in utero* opioid exposure is associated with lower birth weight and adverse neurodevelopmental outcomes in childhood, including cognitive deficits, attention deficit hyperactivity disorder (ADHD), aggression, impaired language development, and decreased social maturity^6–8^. However, large epidemiological studies evaluating long term behavioral outcomes of children exposed to *in utero* opioids have been difficult to perform due to confounding environmental variables including: genetic and epigenetic factors, quality of caregiving, continued parental substance abuse with its impact on the maternal-infant dyad, and other socioeconomic variables which can significantly affect neurodevelopmental outcomes^9^. Consequently, the development of ontogenetic rodent models of opioid exposure is necessary to enable investigation of the biological mechanisms mediating deficits as well as testing alternative treatment avenues for postnatal withdrawal.

To date, there have been a limited number of rodent studies evaluating early-life developmental milestones following *in utero* opioid exposure. Current literature on early developmental effects of *in utero* opioid exposure in pre-clinical models demonstrates decreased birth weight following methadone and buprenorphine exposure^10–12^. Increased latency to right has been observed following *in utero* morphine exposure and is suggestive of different classes of opioids having variable effects on developmental outcomes^13,14^. Opioids exert their pharmacologic effects by activating the endogenous opioid system. While opioids are prescribed for their analgesic effects, acute activation the µ-opioid receptor (MOR) by these medications has also been associated with feelings of euphoria, award reinforcement, and increased socio-emotional processing, which are linked to the drugs’ potential for misuse^15^. Despite the rising incidence of Oxy misuse, there is a paucity of literature evaluating the effects of Oxy, a µ- and κ-agonist, on early developmental behaviors. κ-agonists are of particular interest because over-activation of κ-opioid receptors (KOR) by dynorphin upregulation has been implicated in withdrawal physiology and depressed mood in humans, along with decreased social play in juvenile rodents^15,16^. In addition, most rodent opioid exposure models begin exposure mid-pregnancy, which may explain inconsistently documented or absent developmental changes^17^. To address the above concerns, we are adopting an ontogenetic model in which opioid exposure spans preconception through early offspring development. This new model also enables us to better understand the ontogenetic impact of short- and long-term opioid exposure on early development in the absence of confounding factors present in clinical observational studies. Specifically, we evaluated the effects of *in utero* Oxy exposure on early developmental and behavioral outcomes in male and female offspring of C57BL/6J mouse dams. We implemented a cross-fostering approach that allows us to compare the neurodevelopmental impact of continued postnatal opioid exposure (long-oxy) to neurodevelopmental outcomes of offspring with *in utero* exposure only until birth (short-oxy) by pairing opioid exposed pups with non-oxy exposed dams.

Overall, we observed differences in spectrotemporal features of affective vocalizations and sex-based differences in weight gain trajectories in offspring exposed to Oxy *in utero* (short-oxy). Continued postnatal Oxy exposure (long-oxy) further impacted weight, communicative behavior and sensorimotor reflexes. Our findings suggest that pups with continued postnatal opioid exposure showed worse overall developmental outcomes compared to pups following opioid cessation at birth, which may have implications regarding the safety of continued opioid treatment as mitigation for clinical NAS symptomology.

## MATERIALS and METHODS

### Animals

#### Animal Ethics, Selection and Welfare

All procedures using mice were approved by the Washington University Institutional Care and Use Committee and conducted in accordance with the approved Animal Studies Protocol. C57BL/6J mice (Jackson Laboratory, stock #: 000664) were housed in individually ventilated translucent plastic cages (IVC) measuring 36.2 × 17.1 × 13 cm (Allentown) with corncob bedding and *ad libitum* access to standard lab diet and water. Animals were kept at 12/12 hour light/dark cycle, and room temperature (20-22 °C) and relative humidity (50%) were controlled automatically.

Adult male and female mice were used for breeding cohorts as described below. Sample sizes were determined by power analyses (*f*=0.40, α=0.05, 1-β=0.80). A total of 24 dams were housed in pairs and randomly selected to receive either the Oxy or Vehicle (Veh) treatment infusion. In addition, another set of pair-housed, drug-na’ve dams served as foster dams. Since an inexperienced dam can exhibit poor maternal behavior with her first litter, all females were first bred to an age-matched male at postnatal day (P)60. Following weaning of the first litter, treatment dams underwent surgical subcutaneous pump placement at P95 followed by a one-week recovery period (Figure 1A). Afterwards, each female dam was placed into an individual cage containing a male for breeding. Foster dams were bred at the same time and remained untreated throughout pregnancy. Following 20 days of co-habitation, cages were checked daily for pups. Upon detection, dam and litter were moved to a new cage, without the male, and culled to 6-8 pups per litter with equal males and females when possible. To evaluate the behavioral impact of early opioid cessation in the developing offspring (short-oxy and short-veh), half of the litters were cross-fostered at this time to drug-na’ve dams by removing the pups from their biological dam and transferring them to the nest of the lactating foster dam with two of her own pups of the same approximate age (Figure 1B)^18^. The remaining litters were reared by the biological dam and exposed to prolonged vehicle (long-veh) or oxy (long-oxy) through lactation (Figure 1A,B). To control for litter effects, each group included multiple, independent litters. All mice were weaned at P21 and group-housed by sex with random assignment for drug/dam. Experimenters were all female and blinded to group designations during testing.

**Figure 1.**
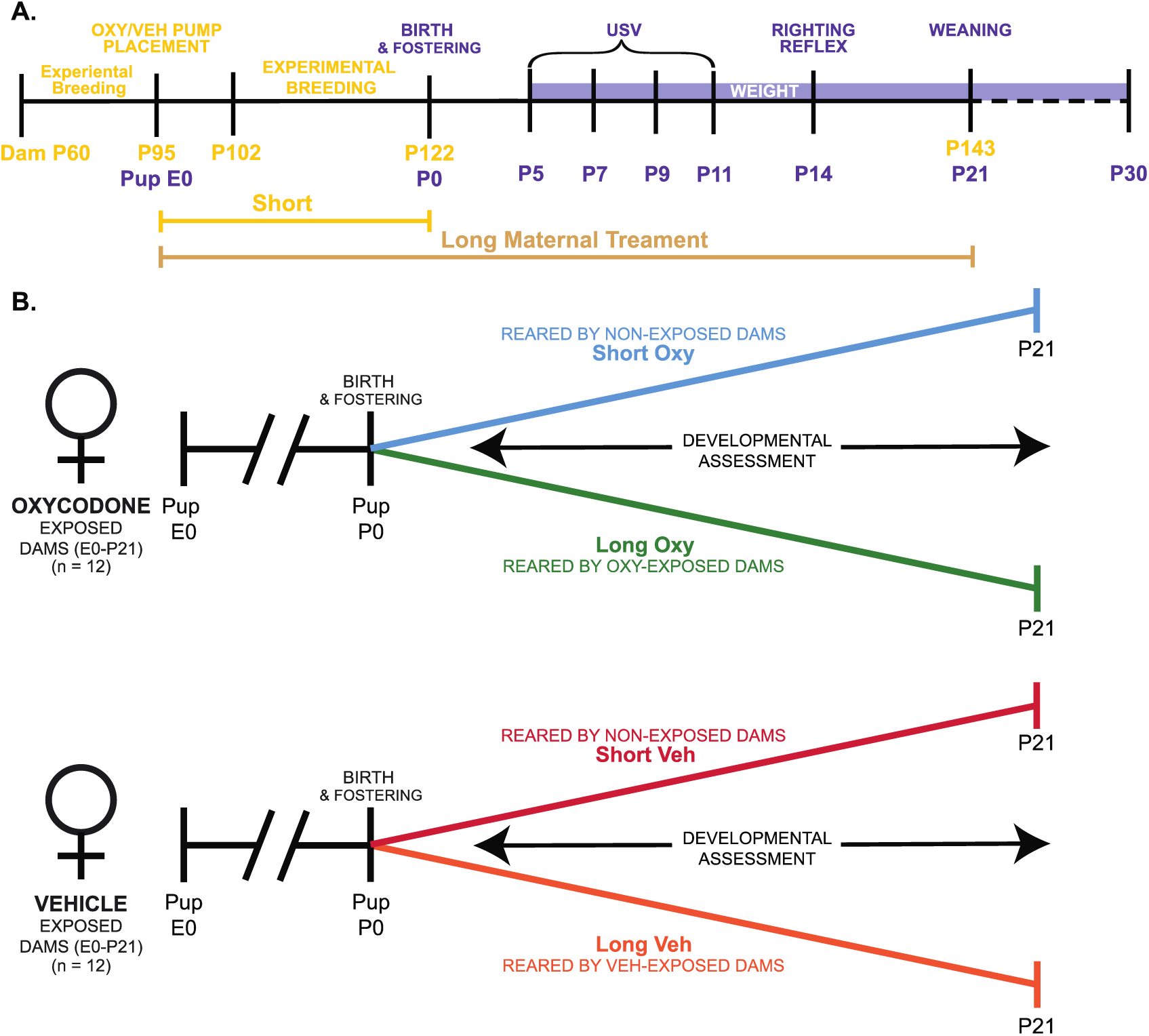
Ontogenetic model of prolonged and limited maternal Oxy exposure. **A**. Schematic of the paradigm for maternal Oxy exposure, including duration of Short and Long maternal treatment, and dam ages at at experimental manipulations in yellow and pup age for behavioral tests in purple. **B**. Outline of the four different experimental groups, including Oxy or Veh exposure, and rearing dam exposure status.

#### Drug Dosage

The dosage of Oxy (Sigma-Aldrich, Saint Louis, MO; Lot#: SLBX4974) administration was guided by previous literature with concentrations ranging from 0.5 mg/kg/day to 33 mg/kg/day^19–21^. Based on this dosage range, we generated our own dosage curve through continuous Oxy administration to pregnant dams at 5, 10, or 15 mg/kg/day using the subcutaneous Alzet 2006 model pump (Durect Corporation, Cupertino, CA; Lot #: 10376-17). We chose the dose of 10 mg/kg/day administered at 0.15 ul/hr to pregnant dams as increased concentrations at 15 mg/kg/day resulted in lower litter success rate (vehicle, 5 & 10 mg/kg/d: 100% success rate; 15 mg/kg/d: 80% success rate).

#### Surgery and Drug Delivery System

Female dams were anesthetized at P95 with isoflurane (5% induction, 2% maintenance, 0.5 l/min) and placed in the mouse adapter (Stoelting, Wood Dale, IL). Body temperature was maintained at 37 °C using a heating pad. The dorsum of the back was shaved and a ∼1 cm horizontal incision was made below the scapulae with subsequent formation of a subcutaneous pocket. The Alzet pump was implanted and continuously infused either Oxy or sterile 0.9% NaCl (Veh) over a period of 60 days. The pump duration allowed for adequate post-surgical recovery time, breeding, and administration of treatment through weaning of offspring at P21. In addition, the use of a subcutaneous pump limited unwanted maternal stress that can occur with daily injections.

### Behavioral Testing

#### Maternal Isolation-Induced Ultrasonic Vocalization Recording

Ultrasonic vocalization (USV) recordings were performed on P5, P7, P9, and P11 (Figure 1A). Dams were removed from the home cage and placed into a clean IVC for the duration of testing. The home cage with the pups in nest was placed into a warming box (Harvard Apparatus) for 10 minutes prior to the start of testing to ensure acclimation of body temperature, as low pup body temperature increases USV production^22^. Skin surface temperature was recorded before placement in the USV recording chamber via a noncontact HDE Infrared Thermometer.

For recording, pups were individually removed from the home cage and placed into an empty standard mouse cage (28.5 × 17.5 × 12 cm) inside a sound-attenuating chamber (Med Associates). USVs were recorded via an Avisoft UltraSoundGate CM16 microphone placed 5 cm away from top of cage, Avisoft UltraSoundGate 116H amplifier, and Avisoft Recorder software (gain = 3 dB, 16 bits, sampling rate = 250 kHz). Pups were recorded for 3 min, after which they were weighed and returned to home cages. Frequency sonograms were prepared from USV recordings in MATLAB [frequency range = 25-120 kHz, Fast Fourier Transform (FFT) size = 512, overlap = 50%, time resolution 1.024 s, frequency resolution = 488.2 Hz]. Individual calls and other spectrotemporal features were identified from the sonograms adapted from validated procedures^23–25^.

#### Developmental Reflexes and Milestones Assessment

Mice were evaluated for achievement of physical and behavioral milestones from early development through early juvenile stage. Weight was measured at 10 time points: P5, P7, P9, P11, P14, P23, P25, P27 and P30. A visual inspection of normal physical milestone attainment was performed with evaluation for detached pinnae at P5 and eye opening at P14. Righting reflex was assessed at P14 as follows: each mouse was placed prone onto its abdomen and quickly pronated 180° to its back in a smooth motion. The time for the mouse to right itself with all four paws positioned underneath the abdomen was recorded ^20^. Each mouse underwent three timed trials, which were averaged for analysis.

### Statistical Analyses

SPSS (IBM, v.25) were used for all statistical analysis. Data was screened for missing values, influential outliers, fit between distributions and the assumptions of normality and homogeneity of variance. Variables that violated assumptions of normality (including number of USV calls, mean pitch, pitch range and peak power) were square root-transformed. Data were analyzed using hierarchical linear models with sex clustered within litters, and age clustered within individual pups. Fixed factors were dam, drug, sex and, where appropriate, age. Age was also treated as a random repeated effect and was grand mean-centered for analysis. As litter size can influence behavior and litter cannot be separated from drug treatment in this study, all models included litter size as a covariate. Probability value for all analyses was *p* < 0.05. Test statistics and analysis details are provided in Table 1. The datasets generated for this study are available upon reasonable request to the corresponding author.

**Table 1.**
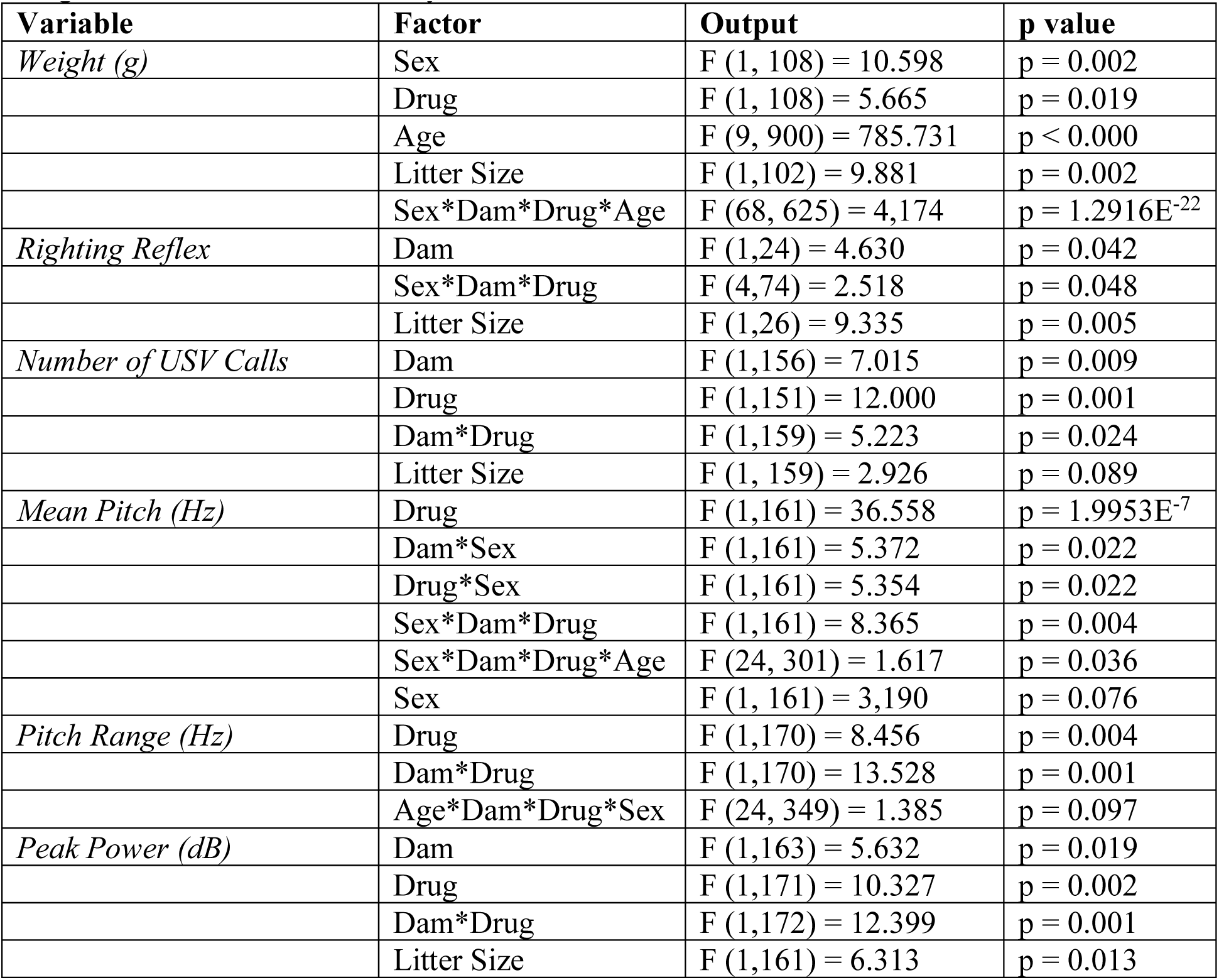
Test statistics from hierarchical linear models. Showing significant main and interaction effects. Age was grand mean-centered for analysis, where included. Litter size was included as a covariate.

## RESULTS

### Oxycodone impacted developmental weight trajectories differentially by sex and exposure duration

We examined effects of Oxy administration on gross and sensorimotor development in mice from birth throughout the early juvenile stage (Figure 2A). To evaluate general health and gross development, we assessed appearance of physical milestones and weight. No differences were observed between groups for pinnae detachment at P5 or eye opening by P14. In our analysis of weight, we found male mice weighed significantly more than females in all groups at all ages, therefore weight data is segregated by sex (Figure 2B, C) from the full factorial linear mixed model including sex, drug, and duration as factors.

**Figure 2.**
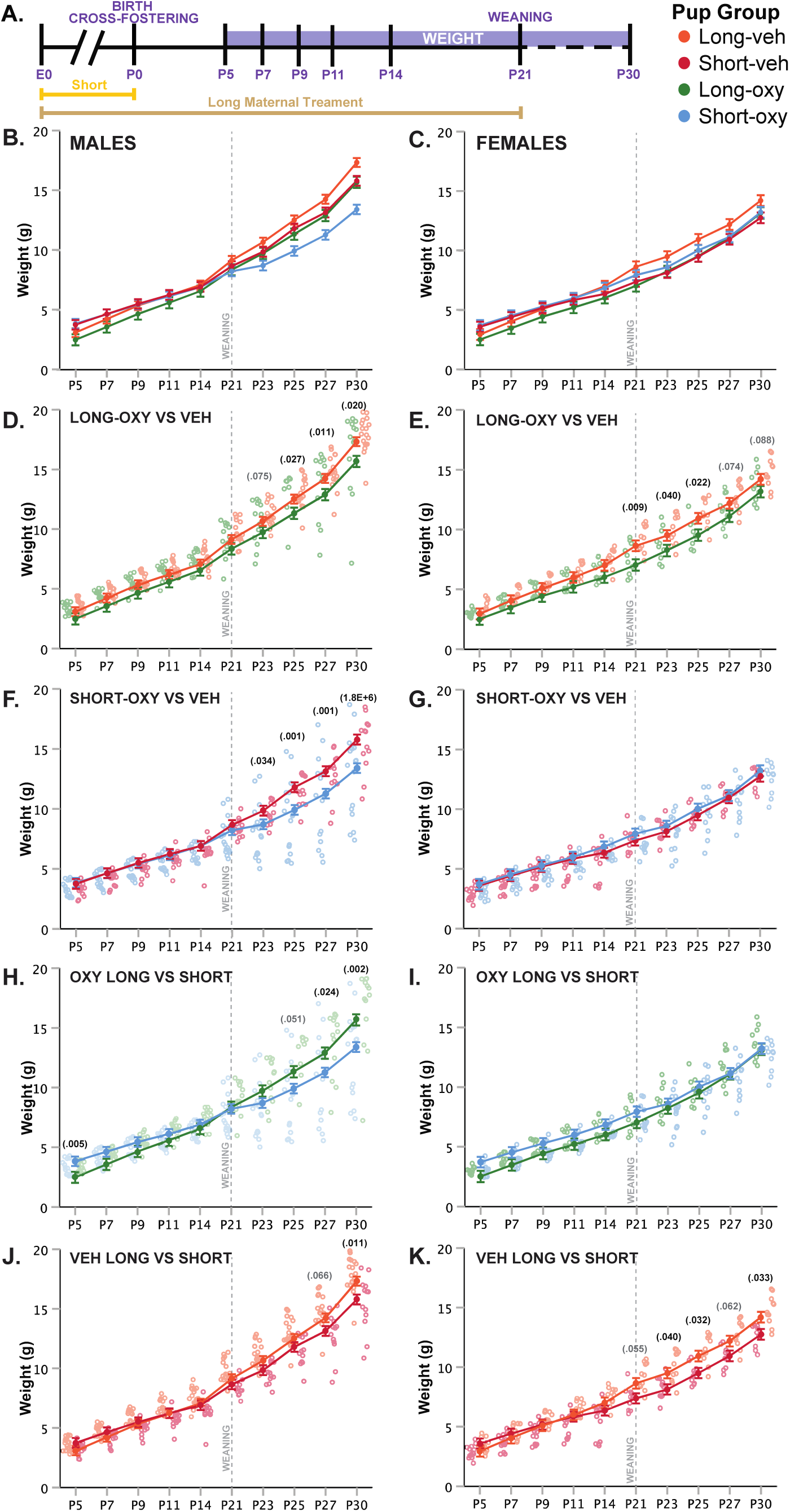
Prolonged and limited Oxy exposure, as well as cross-fostering, decreases weight gain late in development. **A**. Schematic of the treatment paradigm for maternal Oxy exposure and weight measurements throughout development. **B-C**. Line graph of weight in (**B**.) male and (**C**.) female offspring (sex, p=0.002; drug, p=0.019; age, p<0.000; sex × dam × drug × age, p=1.2916E^-22^). **D-E**. Long-oxy exposure, relative to long-veh exposure, led to significantly lower weights post-weaning in (**D**.) male (P23, p = 0.075; P25, p = 0.027; P27, p = 0.011; P30, p = 0.020) and (**E**.) female offspring (P21, p = 0.009; P23, p = 0.040; P25, p = 0.022; P27, p = 0.074; P30, p = 0.088). **F-G**. Short-oxy exposure, relative to short-veh exposure, led to significantly lower weights post-weaning in (**F**.) male offspring only (P23, p = 0.034; P25, p = 0.001; P27, p = 0.001; P30, p = 0.000018), with no effects in (**G**.) female offspring. **H-I**. Short-oxy exposure with cross-fostering, relative to long-oxy exposure, led to decreased weight in adolescence in (**H**.) male pups only (P5, p = 0.005; P25, p = 0.051; P27, p = 0.024; P30, p = 0.002), with no effects in (**I**.) female offspring. **J-K**. Short-veh exposure with cross-fostering, relative to long-veh exposure, led to decreased weight post-weaning in (**J**.) male (P27, p = 0.066; P30, p = 0.011) and (**K**.) female offspring (P21, p = 0.055; P23, p = 0.040; P25, p = 0.032; P27, p = 0.062; P30, p = 0.033). Closed circles depict mean weight, with litter size as a covariate (p = 0.002), while open circles depict individual weights. Gray vertical line indicates date of weaning. * denotes significance at p < 0.05, # indicates trend at p < 0.1.

Long-oxy exposure led to an overall decrease in weight compared to long-veh exposure, which was more pronounced in male offspring. Long-oxy-exposed male offspring exhibited significantly reduced weights compared to long-veh offspring post-weaning at P23, P25, P27 and P30 (Figure 2D). Female long-oxy offspring showed significantly reduced weight compared to long-veh controls at P21, P23, and P25, with non-significant reductions at P27 and P30 (Figure 2E). These data indicate that overall long-oxy exposure reduces weight across development in male and female offspring, with the effect on weight gain compounding once potentially compensatory maternal care is lost post weaning.

We then evaluated the potential effects of early opioid cessation (short-oxy) on weight gain in male and female offspring, and once more found male offspring susceptible to Oxy effects. In contrast to long exposures, an overall reduction in weight was not observed with short-oxy exposure compared to short-veh exposure. However, short-oxy males showed a precipitous decrease in weight gain trajectory after weaning from the foster dam at P23, P25, P27, and P30 as compared to short-veh controls (Figure 2F). Female short-oxy offspring showed no difference in weight across development compared to short-veh controls (Figure 2G). Clinically, male infants are more susceptible to NAS^26^, so the precipitous decrease in weight gain trajectory in the short-oxy male offspring may be associated with withdrawal symptomatology unmasked by cessation of care under a foster dam.

We also examined weight trajectories between short and long vehicle-exposed groups. In vehicle-exposed males, cross-fostering was associated with decreased weights in the short-veh group at P30, with a trend towards decreased weight at P27, relative to the long-veh group (Figure 2J). Of interest, cross-fostered female pups (short-veh) weighed less compared to long-veh female pups post-weaning at P23, P25 and P30 (Figure 2K). These findings suggest cross-fostering alone can influence post-weaning weight trajectories in a sex-dependent manner. However, it is noteworthy that the decrease of weight gain trajectories in the male short-oxy group persisted above and beyond the observed decreased weights in the male short-veh controls, indicating *in utero* Oxy exposure affects weight gain when controlling for cross-fostering (Figure 2B and 2F).

We have shown so far that Oxy exposure, compared to Veh, decreased post-weaning weight following both long and short exposures. We next sought to determine how the duration of Oxy exposure influences weight by assessing the weight gain trajectory differences between long- and short-oxy pups. Comparisons between the male mice showed short-oxy pups initially weighed more than long-oxy pups at P5. However, after the short-oxy male pups were weaned at P21, their weights decreased relative to the long-oxy group at P27 and P30 (Figure 2H). In addition, a subset of the short-oxy mice required saline injections at P23-P25 due to skin tenting, hunched posture, and significant weight loss concerning for dehydration. Following saline injections, recovery was noted in two of the three affected mice with one associated mortality. No differences in weight gain between short and long groups were detected in female Oxy-exposed pups (Figure 2I). Overall, early Oxy cessation was associated with increased weights at very early postnatal ages, followed by weight loss in males at weaning.

### Prolonged oxycodone exposure altered sensorimotor reflex in female offspring only

Righting reflex at P14 was examined as an assessment of sensorimotor milestones, early gross locomotor abilities and general strength (Figure 3A). In males, no difference in latency to right was noted between the long-oxy or long-veh groups (Figure 3B). An increased latency to right was demonstrated in long-oxy male pups compared to short-oxy. However, cross-fostering may have a confounding effect on the righting reflex in male pups, since long-veh males also exhibited an increased latency to right relative to short-veh males. In females, long-oxy pups exhibited significantly increased latency to right relative to long-veh controls (Figure 3C). No significant differences in the righting reflex were observed in the short-oxy or short-veh female offspring. Together, these data indicate that females, but not males, are susceptible to the effects of prolonged developmental Oxy exposure on attainment of the sensorimotor reflex.

**Figure 3.**
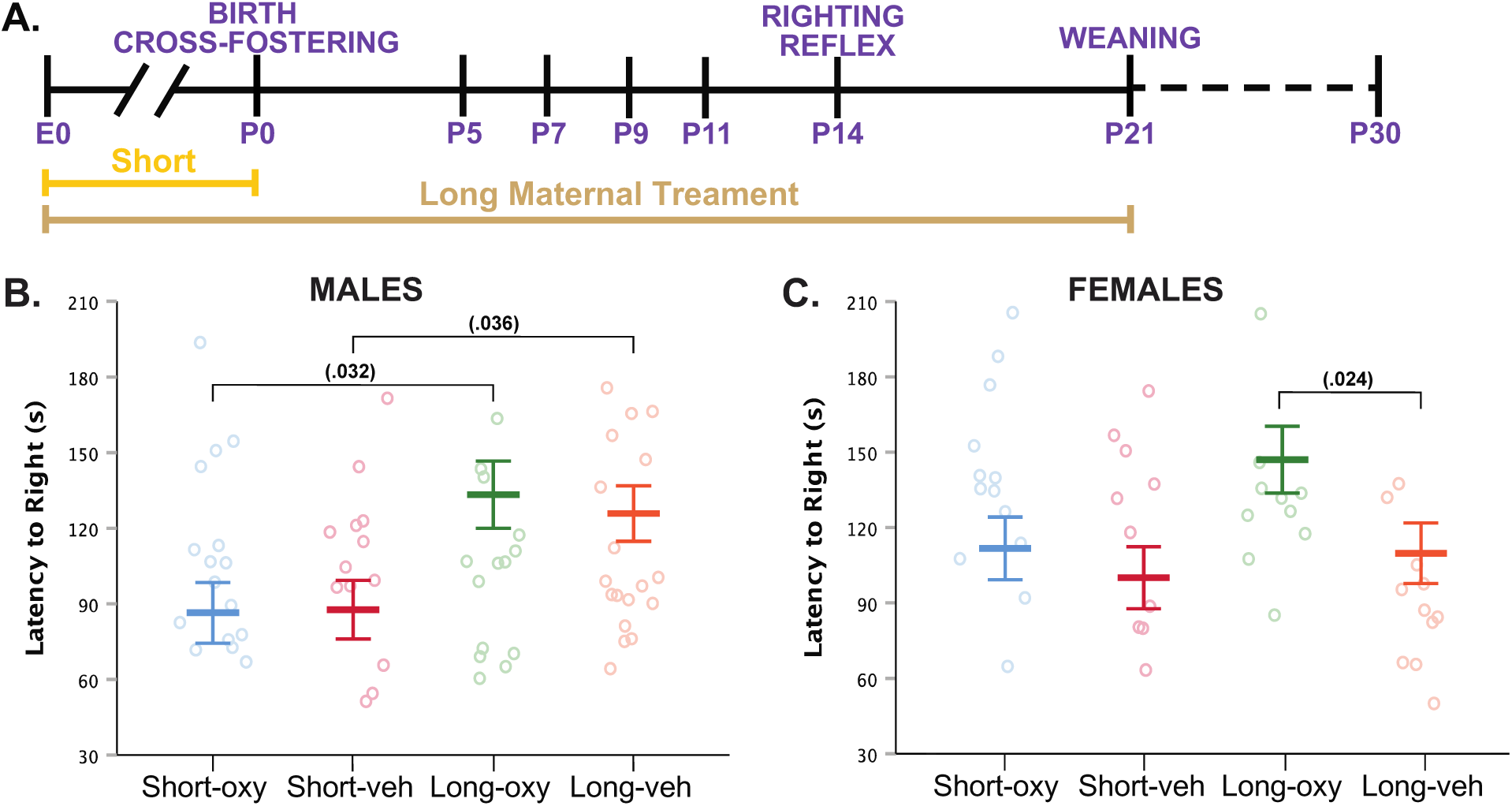
Oxy exposure delays maturing of sensorimotor reflexes. **A**. Schematic of the treatment paradigm for maternal Oxy exposure and righting reflex measurement throughout development. **B-C**. Mean latency to right in (**B**.) male and (**C**.) female offspring (sex × dam × drug × age, p = 0.048; dam, p = 0.042). Long-oxy exposure led to significantly longer latency to right relative to short-oxy (p = 0.032) in (**B**.) male pups and relative to long-veh (p = 0.024) in (**C**.) female pups. Male pups exposed to long-veh also show longer latency to right relative to short-veh (p = 0.036). Closed circles depict mean latencies, with litter size as a covariate (p = 0.005), while open circles depict individual latencies. Error bars represent standard error. * denotes significance at p < 0.05.

### Oxycodone exposure disrupts early communicative behaviors

Language delays have been demonstrated in toddlers with prenatal opioid exposure^7^. Therefore, we assessed early affective and communicative behaviors by evaluating maternal isolation-induced USVs. USVs are an affective and communicative response that elicits maternal search and retrieval, lactation, and caretaking behaviors^23,27^. As a result, characterization of quantity and quality of USV calls has been used in the rodent literature as a model for investigating early communicative deficits^21^. Here, we quantified USV production and spectrotemporal features to examine the influence of Oxy on early communicative behaviors during the first two weeks of life (Figure 4A). If sex had a significant main effect, findings are shown segregated by sex. Overall, we detected a highly significant effect of continued Oxy exposure on USV production. Specifically, long-oxy pups produced significantly fewer USVs relative to long-veh pups and short-oxy pups (Figure 4B), which persisted from P5 through P11 (Figure 4C).

**Figure 4.**
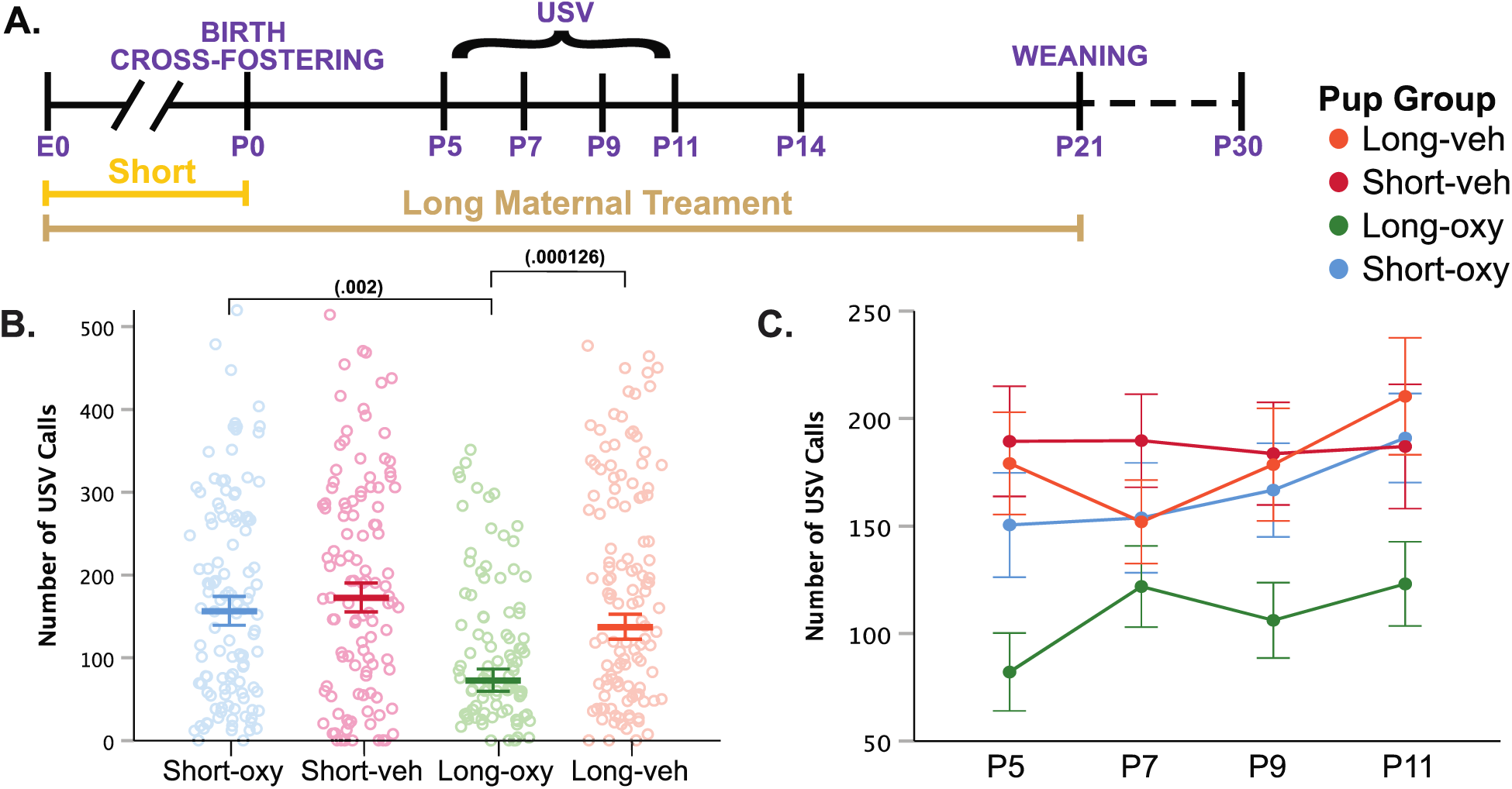
Prolonged Oxy exposure decreases pup USV call production. **A**. Schematic of the treatment paradigm for maternal Oxy exposure and USV measurements throughout development. **B**. Cumulative means number of USV calls (dam, p = 0.009; drug, p = 0.001; dam × drug, p = 0.024). Prolonged Oxy exposure led to decreased number of calls relative to Veh (p = 0.000126) and relative to short-oxy exposure (p = 0.002). **C**. Line graph of mean call number at all time points. Closed circles depict mean call number, with litter size as a covariate, and open circles depict individual call numbers. Error bars represent standard error. * denotes significance at p < 0.05.

Beyond call numbers, spectrotemporal USV features such as duration, pitch frequency, and power (loudness), inform of an affective component to USV characteristics^29^. In previous analyses of USV spectrotemporal features in mouse models of intellectual and developmental disorder risk factors and other early drug exposure models, we and others have demonstrated the vulnerability of these features to genetic and early environmental insults^23,24,30,31^. We examined call features including call duration, pitch range and mean, peak power, and fraction of calls with a pitch jump. Long-oxy administration narrowed the USV pitch range compared to USVs produced by short-oxy pups and long-veh controls (Figure 5A). Long-oxy exposure also led to a highly significant reduction in mean pitch of USVs in long-oxy male pups compared to short-oxy and long-veh males (Figure 5B). Interestingly, USVs produced by long-oxy female pups did not show a significant difference in pitch compared to long-veh females (Figure 5C). However, female short-oxy offspring did exhibit USVs with significantly lower mean pitch relative to short-veh. Thus, *in utero* Oxy exposure was associated with changes in affective components of communication, the significance of which warrants further investigation. Since opioid withdrawal has been associated with high-pitched crying and increased agitation, we also assessed for alterations in USV peak power. Short-oxy exposure resulted in louder USV calls compared to those produced by long-oxy pups and short-veh controls (Figure 5D). The increased peak power, or loudness, in only short-oxy pup calls may be temporally related to onset of withdrawal after drug cessation at P0, relative to the long-oxy pups which are weaned off the drug at P21.

**Figure 5.**
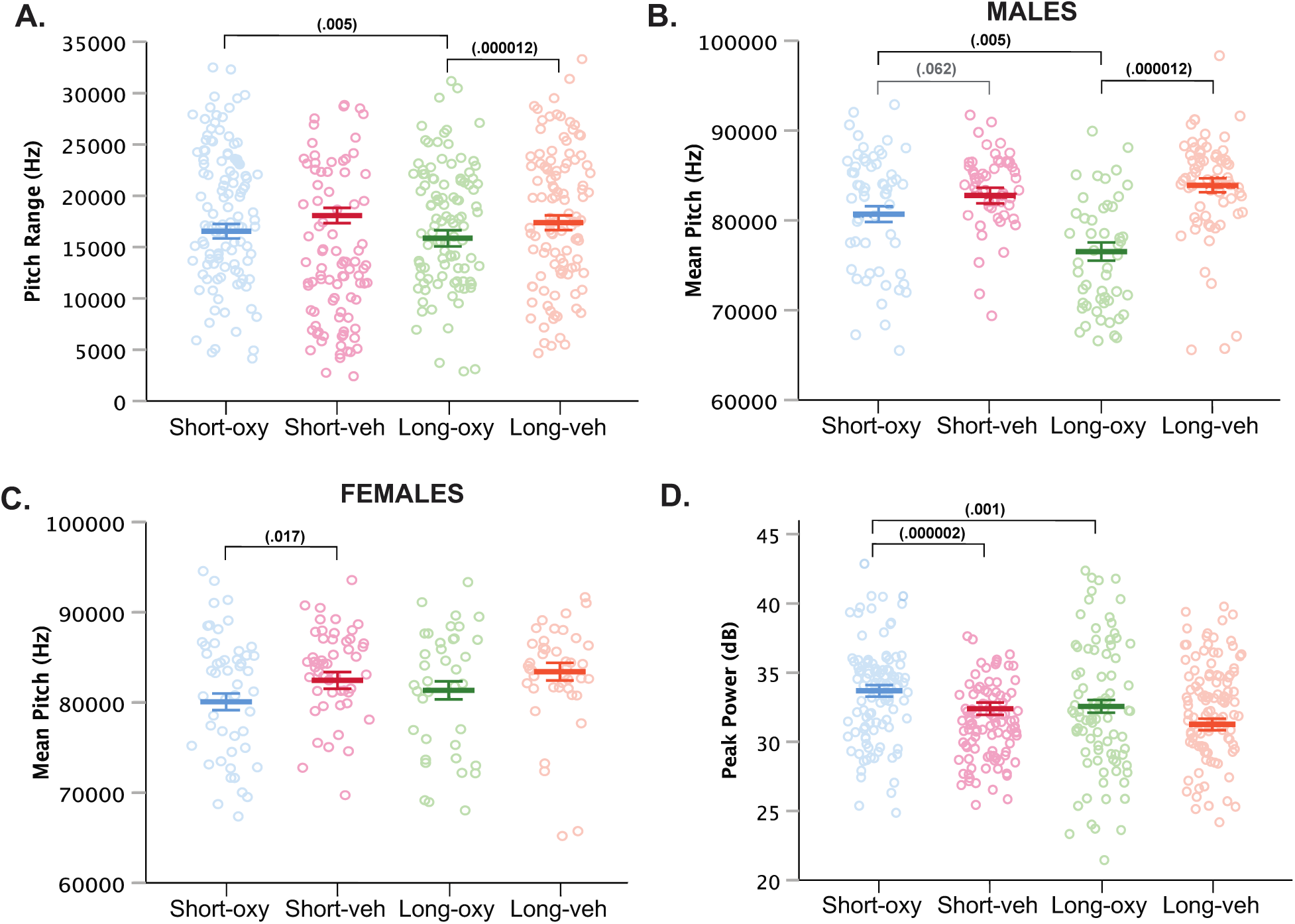
Prolonged and short-oxy exposure affect spectrotemporal features of pup USV calls. **A**. Cumulative means of pitch range (Hz) of USV calls (drug, p = 0.004; dam × drug, p = 0.001; sex × dam × drug × age, p = 0.097). Long-oxy exposure led to lower pitch range relative to Veh (p = 0.000012) and relative to short-oxy exposure (p = 0.005). **B-C**. Cumulative mean pitch (Hz) in (**B**) male and (**C**) female offspring (drug, p = 1.9953E^-7^; sex × dam, p = 0.022; sex × drug, p = 0.022; sex × dam × drug, p = 0.004; sex × dam × drug × age, p = 0.036; sex, p = 0.076). Long-oxy exposure in (**B**.) male pups led to decreased pitch relative to Veh (p = 5.1573E^-9^) and relative to short-oxy (p = 0.016). Short-oxy exposure led to a marginal decrease in pitch relative to Veh (p = 0.062) in (B) male pups and a significant decrease in (**C**) female pups (p = 0.017). **D**. Cumulative means of peak power (dB) (dam, p = 0.032; drug, p = 0.002; dam × drug, p = 0.001). Short-oxy exposure leads to significantly higher peak power relative to long-oxy exposure (p = 0.001) and relative to short-veh (p = 0.000002). Closed circles depict means, with litter size as a covariate, while open circles depict individual measurements. Error bars represent standard error. * denotes significance at p < 0.05, # indicates trend at p < 0.1.

Overall, long-oxy pups demonstrated significant decreases in number of USVs along with a narrower pitch frequency range and mean pitch in male pups. Short-oxy pups produced a similar number of USVs compared to controls, yet those calls were louder than controls and long-oxy calls, and lower in mean pitch when produced by females. Together, our ontogenetic model of *in utero* Oxy exposure demonstrates some alterations in loudness of affective calls, while the prolonged Oxy exposure further shows alterations in number and spectrotemporal features of early communicative and affective behaviors.

## DISCUSSION

Here we present a novel model to investigate the ontogenetic impact of *in utero* only (short) versus continued mitigating opioid exposure (long) on early neurodevelopmental outcomes, while controlling for confounding factors present in clinical observational studies.. *In utero* Oxy exposure decreased weight gain trajectory following weaning from foster dams, in male offspring. Further, *in utero* Oxy-exposed male and female pups showed alterations in the spectrotemporal features of USVs. Meanwhile, offspring with prolonged Oxy exposure until weaning at P21 demonstrated poorer neurodevelopmental outcomes compared to mice exposed only until birth. Notably, continued postnatal Oxy exposure was associated with decreased weight gain trajectory, impaired motor reflexes, and abnormal early communication behaviors. Both male and female offspring in prolonged Oxy exposure groups had decreased weight gain following weaning at P21, with delayed latency to right observed in females. In addition, prolonged Oxy-exposed offspring had significantly reduced USV production and alterations in spectrotemporal features reflecting affective and early communicative impairment.

### Oxycodone exposure impairs attainment of physical and motor development

Decreased fetal growth can be used as a general indicator of harmful *in utero* exposures^2^. The association between birth weight and decreased infant survival is highly robust, though the underlying biological mechanisms are not always clearly understood^32,33^. We did not obtain birth weights at P0 in order to minimize animal handling which can reduce behavioral and hormonal reactivity to stress and confound behavioral testing results^34^. Initial weight assessment occurred at P5 with no observed effect of *in utero* or prolonged Oxy exposure relative to Veh controls. Overall, human literature shows low birth weight in the setting of maternal methadone use during pregnancy, but no evidence of low birth weight following *in utero* exposure to other opioids including codeine, tramadol, hydrocodone, or Oxy^32^. Our finding is consistent with existing Oxy human literature that shows no reported association between Oxy and low birth weight in neonates^32,35^.

Interestingly, female offspring exposed to continued Oxy showed a significant decrease in weight after weaning at P21 (human equivalence of age 2-3 years in terms of brain development)^36–38^, which persisted through the end of experimental testing at P30 (the middle of juvenile development in mice)^38^. Male weights, after prolonged exposure, showed a significant decline on P25-P30 as compared to Veh controls. Given that these offspring were weaned from Oxy postnatally, acute onset of decreased weight gain after separation from the dam at P21 is less likely related to withdrawal symptomatology. Human studies of prenatal opioid exposure have described decreased adaptive behaviors during infancy through toddlerhood^39^. Cessation of care from the biological dam may potentially have uncovered deficiencies in self-care of offspring^6^. Furthermore, maternal environment is a key regulator of offspring development with documented deficits in quality of maternal care in opioid-exposed dams^40,41^. The decreased weight gain post-weaning could also stem from maladaptive feeding behaviors secondary to Oxy exposure, since the opioid system has a strong role driving food intake homeostasis^42^.

Early Oxy cessation significantly decreased weight gain trajectory in a sex-specific manner not observed in the prolonged Oxy exposure cohort. Interestingly, *in utero* exposure led to significantly higher weight at P5, though both groups show comparable averages at P21. Female weights following *in utero* Oxy exposure do not show any significant weight differences from controls. Males exposed to Oxy *in utero* showed a rapid, significant and persistent decrease in weight gain following weaning. Since the *in utero* exposure group was cross-fostered at birth to a non-drug exposed dam, normal weight gain trajectory was potentially maintained through adequate maternal care. A further explanation could be the “two-hit” hypothesis in which early-life susceptibility, such as abstinence and withdrawal following *in utero* opioid exposure, compounded with the postnatal stress of weaning, precipitated a weight loss phenotype^43,44^. Perhaps, males are more sensitive to early life stressors and may have long-term consequences from *in utero* opioid exposure compared to females. Male human neonates are more at risk for developing NAS compared to females, so perhaps long-term changes in opioid circuitry governing feeding behaviors could explain the abnormal weight trajectory in male mouse offspring post-weaning^26^. Prospective human studies evaluating the long-term effects of *in utero* opioid and effects on weight trajectory during childhood through adulthood have not been performed to our knowledge. The altered weight trajectory findings in both the short-Oxy and long-Oxy suggest a potential role of *in utero* opioid exposure on long-term impact on growth that requires further evaluation in the human literature.

In the vehicle-treated groups, cross-fostered pups showed decreased weight gain trajectory after weaning relative to pups reared by a biological dam. Our observation of decreased weight gain following weaning in Veh-exposed cross-fostered pups may be related to potential alterations in emotionality and stress responses secondary to confounders involved with cross-fostering, such as early handling^34^. Regardless, the effect of cross-fostering on weight did not mask our ability to identify effects of *in utero* Oxy exposure on weight in males. Indeed, the effect of *in utero* Oxy exposure on weight occurred at additional younger ages and with a larger magnitude than *in utero* Veh exposure and persisted when controlling for effects of litter and cross-fostered dam status. Similarly, in females, weight reduction was observed following prolonged Oxy and *in utero* Oxy exposure, and *in utero* Veh exposure compared to the prolonged Veh exposure control group. Interestingly, as discussed below, cross-fostering resulted in a shorter latency to exhibit the righting reflex compared to the pups reared by biological dams. This finding indicates that cross-fostering did not impair the development of sensorimotor reflexes. Therefore, despite the independent effect on weight by cross-fostering, this method was valuable in allowing us to cease Oxy exposure at birth and thus observe effect of Oxy limited to *in utero* development. Furthermore, these findings highlight the importance of including proper cross-foster control groups in study designs for accurate interpretation of results.

Prenatal opioid exposure has also been associated with delays in attainment of motor milestones in children. A meta-analysis by Yeoh *et. al* (2019) detected significant delays in motor outcomes in children aged 0-6 years that experienced prenatal opioid exposure^45^. We assessed the righting reflex at P14, the beginning of the visual critical period and an age at which mice should be fully ambulatory^46–48^. The righting reflex corrects the orientation of the body from an off axis position^49^. Proper execution of the reflex requires a combination of visual, vestibular, and somatosensory system inputs to make appropriate postural adjustments through neural pathways within the brain and cerebellum. Interestingly, females in the prolonged Oxy exposure group had significantly increased latency to right compared to Veh controls (Figure 4B). There was no significant difference in righting reflex latency between females exposed to Oxy and Veh *in utero* (Figure 4B). Hence, only continued postnatal Oxy exposure seems to result in delayed sensorimotor development. Previously, increased latency to right has been demonstrated with *in utero* morphine exposure in both male and female rat pups, but rodent studies evaluating effects of opioids on the righting reflex have been limited^13^. Though the exact mechanism of action is unclear, significant evidence in the literature demonstrates selective vulnerability of cerebellar granule neuroblasts to opioids, along with opioids’ negative effects on neuronal somatosensory cortex development^50,51^. Multiple studies have further linked opioid exposure to increased apoptosis and decreased differentiation of Purkinje cells in the cerebellum^50^. Additionally, perinatal morphine treatment in rats decreased total number of neurons in the somatosensory cortex^51^. Thus, it is possible multiple circuits are mediating the effect. Overall, the sex-specificity of the effect is also interesting. This observed sex bias could be related to sex-specific dimorphic alterations in catecholamine levels in the cerebellum as shown in a previous study of prenatal morphine exposure^52^. However, future studies will be necessary to delineate the mechanisms underlying this behavioral phenotype, and its sex-specific expression.

### Ontogenetic oxycodone exposure may delay early communicative behaviors and alter spectrotemporal features of USV

Currently, studies exploring the effects of prenatal opioid exposure on language development in children demonstrate equivocal results^39^. Previous work has identified language development impairments following *in utero* exposure to methadone or heroin^39^. However, several of these analyses did not control for important confounders such as socioeconomic status or maternal use of other substances. In general, large epidemiological studies evaluating impact of prenatal opioid exposure on language development have been difficult to perform due to various confounding environmental variables including quality of caregiving, parental education level, and socioeconomic factors. Despite the advantage of limiting confounds through the use of rodent models, and indications that communicative circuits are conserved between rodents and humans^53^, there have been minimal rodent studies evaluating the effects of prenatal opioid exposure on early communicative behaviors to date. Isolation-induced USVs are a strongly conserved adaptive response of young rodent pups to elicit maternal caregiving responses^27^. Our observed collective decrease in mean USV production following prolonged Oxy exposure, compared to *in utero* exposure and Veh controls, is suggestive of impaired early communication. Previous studies have shown evidence for neuropharmacological modulation of USVs through alteration of mood or arousal state^23,54,55,55^. In addition to serving as an analgesic, Oxy is also a sedative and an anxiolytic agent, which may decrease USV calling in the prolonged Oxy exposure pups by actively suppressing USV circuitry secondary to reduced reactivity to surrounding environmental stressors and decreased respiratory rate^56^. Furthermore, the interplay between pup communication and maternal care is complicated. In vocally impaired pups, decreased USV production has been shown to result in maternal neglect, because without calls the dams cannot locate the pups outside of the nest^57^. Thus, maternal care may be reduced in response to decreased USVs calling by prolonged Oxy-exposed pups. Maternal dam care is also attenuated following opioid exposure during pregnancy, with studies reporting increased time to pup retrieval, decreased nursing and cleaning of pups, and increased maternal self-care time^41^. This reduced maternal care could further disrupt neurodevelopment of the pup, and thus be a possible indirect influence on later adult behaviors. Notably, the *in utero* Oxy offspring demonstrated comparable USV means to Veh controls. The normal USV call production in the *in utero* Oxy exposed group by P5 suggests Oxy reduces USV production in the prolonged exposure groups by acute suppression of USV circuits. Finally, the adequate weight trajectories in the Oxy group further indicate appropriate dam care. Hence, we would hypothesize that maternal care likely did not result in the behavioral deficits observed. Still, to our knowledge, the direct impact of Oxy exposure on maternal behaviors has not been examined and warrants an individual study to assess reciprocity of interactions between pup and dam. Overall, deficits in observed USV production could be the result of a combination of factors, including acute drug effects on circuits, influence of dam care, and opioid-mediated effects on neural development and communication.

Affective characteristics of USVs in rodents are generally thought to communicate different emotional states, such as aggression or pain. Rodent USVs are particularly interesting as they occur only in salient situations such as exposure to painful stimuli, maternal behavior, sexual behavior, or aggression. As has been well-characterized in rats, affective features of rodent USVs may be reflected by alterations in duration, pitch, frequency, and loudness (dB) of the calls^54,58^. In our developmental cohort, pups exposed to prolonged Oxy demonstrated decreased mean pitch (in males only), and narrower pitch range. Interestingly, a previous study administered morphine to adult rats and observed decreased USV pitch, duration, and rate^58^. The decreased pitch and pitch range could be a result of Oxy’s depressive effects on respiration, or of Oxy’s anxiolytic drug properties which may dampen USV circuity. Meanwhile, pups exposed to Oxy *in utero* demonstrated decreased mean pitch of USV calls (females only) and decreased mean pitch range. Of interest, mean peak power, or loudness, was increased in the *in utero* Oxy exposure group compared to prolonged Oxy exposure Veh controls, which could be related to agitation associated with withdrawal. Neonates with NAS are frequently described as exhibiting a high-pitched cry with prolonged periods of inconsolability and crying. Although we did not notice an increase in pitch or frequency within the *in utero* Oxy exposed offspring as would be expected with NAS, the increased power of the USV calls may be interpreted as an affective feature of distress.

Our novel Oxy administration paradigm enables future exploration of withdrawal periods, to determine if NAS following Oxy cessation can be appropriately modeled in rodents. If so, testing of novel agents for treatment of withdrawal symptoms will be possible, with the goal of limiting continued postnatal opioid exposure given the potential long-term side effects of early-postnatal opioid administration on neurodevelopment^59^. Finally, few mouse models of *in utero* opioid exposure currently exist, with the majority of the literature utilizing rat perinatal opioid models. Thus, creation of a mouse model will facilitate genetic manipulations using the established cutting-edge genetic tools available in the mouse to broaden understanding of the mechanisms mediating consequences of early opioid exposure on neurodevelopment.

## Author Contributions

EM contributed to conceptualization, methodology, investigation, data curation, writing (original draft preparation and editing), and funding acquisition for this project.

SS contributed to methodology, validation, formal analysis, writing (original draft preparation and editing), and visualization for this project.

JD contributed to conceptualization, methodology, validation, resources, writing (draft editing), supervision, project administration, and funding acquisition for this project.

RA contributed to conceptualization, methodology, validation, writing (draft editing), supervision, project administration, and funding acquisition for this project.

SEM contributed to conceptualization, methodology, data curation, resources, writing (draft editing), validation, supervision, project administration, and funding acquisition for this project.

## REFERENCES

1. Affairs (ASPA) AS of P. What is the U.S. Opioid Epidemic? HHS.gov. Published December 4, 2017. Accessed June 22, 2020.https://www.hhs.gov/opioids/about-the-epidemic/index.html

2. Haight SC. Opioid Use Disorder Documented at Delivery Hospitalization — United States, 1999–2014. MMWR Morb Mortal Wkly Rep. 2018;67. doi:10.15585/mmwr.mm6731a1

3. Kenan K, Mack K, Paulozzi L. Trends in prescriptions for oxycodone and other commonly used opioids in the United States, 2000–2010. Open Med. 2012;6(2):e41–e47.

4. Wiles JR, Isemann B, Ward LP, Vinks AA, Akinbi H. Current Management of Neonatal Abstinence Syndrome Secondary to Intrauterine Opioid Exposure. J Pediatr. 2014;165(3):440. doi:10.1016/j.jpeds.2014.05.010

5. Hudak ML, Tan RC, Drugs TCO, Newborn TC on FA. Neonatal Drug Withdrawal. Pediatrics. 2012;129(2):e540–e560. doi:10.1542/peds.2011-3212

6. Azuine RE, Ji Y, Chang H-Y, et al. Prenatal Risk Factors and Perinatal and Postnatal Outcomes Associated With Maternal Opioid Exposure in an Urban, Low-Income, Multiethnic US Population. JAMA Netw Open. 2019;2(6):e196405–e196405. doi:10.1001/jamanetworkopen.2019.6405

7. Conradt E, Flannery T, Aschner JL, et al. Prenatal Opioid Exposure: Neurodevelopmental Consequences and Future Research Priorities. Pediatrics. 2019;144(3). doi:10.1542/peds.2019-0128

8. Hunt RW, Tzioumi D, Collins E, Jeffery HE. Adverse neurodevelopmental outcome of infants exposed to opiate in-utero. Early Hum Dev. 2008;84(1):29–35. doi:10.1016/j.earlhumdev.2007.01.013

9. Lutz P-E, Kieffer BL. Opioid receptors: distinct roles in mood disorders. Trends Neurosci. 2013;36(3):195–206. doi:10.1016/j.tins.2012.11.002

10. Kunko PM, Smith JA, Wallace MJ, Maher JR, Saady JJ, Robinson SE. Perinatal methadone exposure produces physical dependence and altered behavioral development in the rat. J Pharmacol Exp Ther. 1996;277(3):1344–1351.

11. Hung C-J, Wu C-C, Chen W-Y, et al. Depression-Like Effect of Prenatal Buprenorphine Exposure in Rats. PLoS ONE. 2013;8(12). doi:10.1371/journal.pone.0082262

12. Chiang Y-C, Ye L-C, Hsu K-Y, et al. Beneficial effects of co-treatment with dextromethorphan on prenatally methadone-exposed offspring. J Biomed Sci. 2015;22:19. doi:10.1186/s12929-015-0126-2

13. Slamberová R, Riley MA, Vathy I. Cross-generational effect of prenatal morphine exposure on neurobehavioral development of rat pups. Physiol Res. 2005;54(6):655–660.

14. Niu L, Cao B, Zhu H, et al. Impaired in vivo synaptic plasticity in dentate gyrus and spatial memory in juvenile rats induced by prenatal morphine exposure. Hippocampus. 2009;19(7):649–657. doi:10.1002/hipo.20540

15. Vanderschuren LJ, Niesink RJ, Spruijt BM, Van Ree JM. Mu-and kappa-opioid receptor-mediated opioid effects on social play in juvenile rats. Eur J Pharmacol. 1995;276(3):257–266. doi:10.1016/0014-2999(95)00040-r

16. Li W, Sun H, Chen H, et al. Major Depressive Disorder and Kappa Opioid Receptor Antagonists. Transl Perioper Pain Med. 2016;1(2):4–16.

17. Richardson KA, Yohay A-LJ, Gauda EB, McLemore GL. Neonatal Animal Models of Opiate Withdrawal. ILAR J. 2006;47(1):39–48. doi:10.1093/ilar.47.1.39

18. Lohmiller J, Swing S.Reproduction and Breeding. In: The Laboratory Rat.; 2006:147–164. doi:10.1016/B978-012074903-4/50009-1

19. Sithisarn T, Legan SJ, Westgate PM, et al. The Effects of Perinatal Oxycodone Exposure on Behavioral Outcome in a Rodent Model. Front Pediatr. 2017;5:180. doi:10.3389/fped.2017.00180

20. Zanni G, Robinson-Drummer PA, Dougher AA, et al. Maternal Continuous Oral Oxycodone Self-Administration Alters Pup Affective/Social Communication but not Spatial Learning or Sensory-Motor Function. bioRxiv. Published online April 5, 2020:2020.04.04.022533. doi:10.1101/2020.04.04.022533

21. Enga RM, Jackson A, Damaj MI, Beardsley PM. Oxycodone physical dependence and its oral self-administration in C57BL/6J mice. Eur J Pharmacol. 2016;789:75–80. doi:10.1016/j.ejphar.2016.07.006

22. Branchi I, Santucci D, Alleva E. Ultrasonic vocalisation emitted by infant rodents: a tool for assessment of neurobehavioural development. Behav Brain Res. 2001;125(1):49–56. doi:10.1016/S0166-4328(01)00277-7

23. Maloney SE, Akula S, Rieger MA, et al. Examining the Reversibility of Long-Term Behavioral Disruptions in Progeny of Maternal SSRI Exposure. eNeuro. 2018;5(4). doi:10.1523/ENEURO.0120-18.2018

24. Maloney SE, Chandler KC, Anastasaki C, Rieger MA, Gutmann DH, Dougherty JD. Characterization of early communicative behavior in mouse models of neurofibromatosis type 1. Autism Res Off J Int Soc Autism Res. 2018;11(1):44–58. doi:10.1002/aur.1853

25. Holy TE, Guo Z. Ultrasonic songs of male mice. PLoS Biol. 2005;3(12):e386. doi:10.1371/journal.pbio.0030386

26. Charles MK, Cooper WO, Jansson LM, Dudley J, Slaughter JC, Patrick SW. Male Sex Associated With Increased Risk of Neonatal Abstinence Syndrome. Hosp Pediatr. 2017;7(6):328–334. doi:10.1542/hpeds.2016-0218

27. Haack B, Markl H, Ehret G. Sound communication between parents and offspring. In:; 2009.doi:10.18725/OPARU-1174

28. Hofer MA, Shair HN, Brunelli SA. Ultrasonic vocalizations in rat and mouse pups. Curr Protoc Neurosci. 2002;Chapter 8:Unit 8.14. doi:10.1002/0471142301.ns0814s17

29. Wöhr M, Schwarting RKW. Affective communication in rodents: ultrasonic vocalizations as a tool for research on emotion and motivation. Cell Tissue Res. 2013;354(1):81–97. doi:10.1007/s00441-013-1607-9

30. Kopp N, McCullough K, Maloney SE, Dougherty JD. Gtf2i and Gtf2ird1 mutation do not account for the full phenotypic effect of the Williams syndrome critical region in mouse models. Hum Mol Genet. 2019;28(20):3443–3465. doi:10.1093/hmg/ddz176

31. Dougherty JD, Maloney SE, Wozniak DF, et al. The disruption of Celf6, a gene identified by translational profiling of serotonergic neurons, results in autism-related behaviors. J Neurosci Off J Soc Neurosci. 2013;33(7):2732–2753. doi:10.1523/JNEUROSCI.4762-12.2013

32. Yazdy MM, Desai RJ, Brogly SB. Prescription Opioids in Pregnancy and Birth Outcomes: A Review of the Literature. J Pediatr Genet. 2015;4(2):56–70. doi:10.1055/s-0035-1556740

33. Basso O, Wilcox AJ, Weinberg CR. Birth Weight and Mortality: Causality or Confounding? Am J Epidemiol. 2006;164(4):303–311. doi:10.1093/aje/kwj237

34. Luchetti A, Oddi D, Lampis V, et al. Early handling and repeated cross-fostering have opposite effect on mouse emotionality. Front Behav Neurosci. 2015;9. doi:10.3389/fnbeh.2015.00093

35. Kelly L, Dooley J, Cromarty H, et al. Narcotic-exposed neonates in a First Nations population in northwestern Ontario: incidence and implications. Can Fam Physician Med Fam Can. 2011;57(11):e441–447.

36. Dobbing J, Sands J. Comparative aspects of the brain growth spurt. Early Hum Dev. 1979;3(1):79–83. doi:10.1016/0378-3782(79)90022-7

37. Workman AD, Charvet CJ, Clancy B, Darlington RB, Finlay BL. Modeling Transformations of Neurodevelopmental Sequences across Mammalian Species. J Neurosci. 2013;33(17):7368–7383. doi:10.1523/JNEUROSCI.5746-12.2013

38. McCarthy MM, Wright CL. Convergence of Sex Differences and the Neuroimmune System in Autism Spectrum Disorder. Biol Psychiatry. 2017;81(5):402–410. doi:10.1016/j.biopsych.2016.10.004

39. Conradt E, Flannery T, Aschner JL, et al. Prenatal Opioid Exposure: Neurodevelopmental Consequences and Future Research Priorities. Pediatrics. 2019;144(3). doi:10.1542/peds.2019-0128

40. Song L, Johnson MD, Tamashiro KLK. Maternal and Epigenetic Factors That Influence Food Intake and Energy Balance in Offspring. In: Harris RBS, ed. Appetite and Food Intake: Central Control. 2nd ed. CRC Press/Taylor & Francis; 2017. Accessed June 23, 2020. http://www.ncbi.nlm.nih.gov/books/NBK453152/

41. Slamberová R, Szilágyi B, Vathy I. Repeated morphine administration during pregnancy attenuates maternal behavior. Psychoneuroendocrinology. 2001;26(6):565–576. doi:10.1016/s0306-4530(01)00012-9

42. Valbrun LP, Zvonarev V. The Opioid System and Food Intake: Use of Opiate Antagonists in Treatment of Binge Eating Disorder and Abnormal Eating Behavior. J Clin Med Res. 2020;12(2):41–63. doi:10.14740/jocmr4066

43. Peña CJ, Nestler EJ, Bagot RC. Environmental Programming of Susceptibility and Resilience to Stress in Adulthood in Male Mice. Front Behav Neurosci. 2019;13. doi:10.3389/fnbeh.2019.00040

44. Nederhof E, Schmidt MV. Mismatch or cumulative stress: toward an integrated hypothesis of programming effects. Physiol Behav. 2012;106(5):691–700. doi:10.1016/j.physbeh.2011.12.008

45. Yeoh SL, Eastwood J, Wright IM, et al. Cognitive and Motor Outcomes of Children With Prenatal Opioid Exposure: A Systematic Review and Meta-analysis. JAMA Netw Open. 2019;2(7):e197025–e197025. doi:10.1001/jamanetworkopen.2019.7025

46. Hooks BM, Chen C. Critical Periods in the Visual System: Changing Views for a Model of Experience-Dependent Plasticity. Neuron. 2007;56(2):312–326. doi:10.1016/j.neuron.2007.10.003

47. Williams E, Scott J. The Development of Social Behavior Patterns in the Mouse, in Relation To Natural Periods 1). Published online 1954.doi:10.1163/156853954X00031

48. Feather-Schussler DN, Ferguson TS. A Battery of Motor Tests in a Neonatal Mouse Model of Cerebral Palsy. J Vis Exp JoVE. 2016;(117). doi:10.3791/53569

49. Righting reflex. In: Wikipedia.; 2020. Accessed June 26, 2020.https://en.wikipedia.org/w/index.php?title=Righting_reflex&oldid=943956135

50. Hauser KF, Khurdayan VK, Goody RJ, Nath A, Saria A, Pauly JR. Selective vulnerability of cerebellar granule neuroblasts and their progeny to drugs with abuse liability. Cerebellum Lond Engl. 2003;2(3):184–195. doi:10.1080/14734220310016132

51. Seatriz JV, Hammer RP. Effects of opiates on neuronal development in the rat cerebral cortex. Brain Res Bull. 1993;30(5):523–527. doi:10.1016/0361-9230(93)90078-P

52. Vathy I, Rimanoczy A, Eaton RC, Katay L. Sex dimorphic alterations in postnatal brain catecholamines after gestational morphine. Brain Res Bull. 1995;36(2):185–193. doi:10.1016/0361-9230(94)00192-4

53. Arriaga G, Zhou EP, Jarvis ED. Of Mice, Birds, and Men: The Mouse Ultrasonic Song System Has Some Features Similar to Humans and Song-Learning Birds. PLOS ONE. 2012;7(10):e46610. doi:10.1371/journal.pone.0046610

54. Vivian JA, Miczek KA. Diazepam and gepirone selectively attenuate either 20-32 or 32-64 kHz ultrasonic vocalizations during aggressive encounters. Psychopharmacology (Berl). 1993;112(1):66–73. doi:10.1007/BF02247364

55. Vivian JA, Miczek KA. Ultrasounds during morphine withdrawal in rats. Psychopharmacology (Berl). 1991;104(2):187–193. doi:10.1007/BF02244177

56. Rao R, Desai NS. OxyContin and Neonatal Abstinence Syndrome. J Perinatol. 2002;22(4):324–325. doi:10.1038/sj.jp.7210744

57. Hernandez-Miranda LR, Ruffault P-L, Bouvier JC, et al. Genetic identification of a hindbrain nucleus essential for innate vocalization. Proc Natl Acad Sci U S A. 2017;114(30):8095–8100. doi:10.1073/pnas.1702893114

58. Vivian JA, Miczek KA. Morphine attenuates ultrasonic vocalization during agonistic encounters in adult male rats. Psychopharmacology (Berl). 1993;111(3):367–375. doi:10.1007/BF02244954

59. Attarian S, Tran LC, Moore A, Stanton G, Meyer E, Moore RP. The Neurodevelopmental Impact of Neonatal Morphine Administration. Brain Sci. 2014;4(2):321–334. doi:10.3390/brainsci4020321

